# Estimating real snakebite incidence in Colombia by using mathematical modelling and statistical inference

**DOI:** 10.1101/2021.12.09.472006

**Authors:** Carlos Bravo-Vega, Camila Renjifo-Ibañez, Mauricio Santos-Vega, Leonardo Jose Leon Nuñez, Teddy Angarita-Sierra, Juan Manuel Cordovez

## Abstract

Snakebite envenoming is a Neglected Tropical Disease affecting mainly deprived populations. Its burden is normally underestimated because patients prefer to seek for traditional medicine. Thus, applying strategies to optimize disease’ management and treatment delivery is difficult. We propose a framework to estimate snakebite incidence at a fine political scale based on available data, testing it in Colombia. First, we produced snakebite fine-scale risk maps based on the most medically important venomous snake species (*Bothrops asper* and *B. atrox*). We validated them with reported data in the country. Then, we proposed a generalized mixed effect model that estimates total incidence based on produced risk maps, poverty indexes, and an accessibility score that reflects the struggle to reach a medical center. Finally, we calibrated our model with national snakebite reported data from 2010 to 2019 using a Markov chain Monte Carlo (MCMC) algorithm and estimated underreporting based on the total incidence estimation. Our results suggest that 10.3% of total snakebite cases are not reported in Colombia and do not seek medical attention. The Orinoco and Amazonian regions (east of Colombia) share a high snakebite risk with a high underreporting. Our work highlights the importance of multidisciplinary approaches to face snakebite.

## INTRODUCTION

Snakebite envenoming is a Neglected Tropical Disease (NTD) characterized by its high mortality and morbidity rates (1–4). The most effective treatment for this disease is the administration of antivenom, which is composed of a mixture of antibodies from one (monospecific) to several (polyspecific) venomous snake species, neutralizing the toxic effects of the venom (5). Although antivenom is life-saving, most patients affected by a snakebite live in remote rural areas where it is not readily available (6,7). Envenomation episodes in rural areas often result in patients seeking traditional healers instead of antivenom therapy (4,8–11). This situation poses a challenge in data collection, which usually does not account for the number of envenoming cases (12,13). Under these circumstances, mapping vulnerable areas become an urgent need because incomplete datasets can overlook the increasing snakebite real burden. As a global effort to reduce snakebite mortality by 50% in 2030, the world health organization proposed a global strategy where one aim corresponds with strengthening surveillance systems, improving antivenom availability, and including this NTD into public health scope. Indeed, the first step to fulfill this objective is to generate reliable estimates of snakebite burden, where disease eco-epidemiology can help pinpoint overlooked vulnerable populations (14,15).

Estimating disease burden by performing cross-sectional studies requires resources that are not readily available(16–18). Therefore, it is necessary to study the natural history of the species involved in the disease and then perform mathematical modeling to estimate disease’ epidemiological parameters (19–22). This biological knowledge becomes critical in megadiverse countries like Colombia, where approximately 319 species of snakes can be found, but only 52 represent a potential risk for humans. Out of these 52 species, 21 belong to the *Viperidae* family and 31 belong to the *Elapidae* family (23–25). These venomous snakes are distributed in most of the territory, and human–snake encounters occur frequently (26–28). The most medically important venomous snake species in the country are *Bothrops asper* and *B. atrox* (Common names: Talla X and Cuatro narices respectively), which are responsible for most of the envenomings (some studies suggest that it could be as much as 80%) (25,28–33). Previous studies for Colombia have proposed that the species range for *B. asper* is the Caribbean, Pacific coast, and the inter-Andean valleys. On the contrary, the species range of *B. atrox* ranges from the Orinoco and Amazonian regions (25). However, their distribution and association with environmental variables are largely absent in scientific studies (34,35).

Snakebite envenoming usually lacks reliable and long-term data accounting for the number of cases in each region in a country. In Colombia, snakebite cases treated in medical centers are reported to Sistema Nacional de Vigilancia en Salud Pública (SIVIGILA), which monitors and collects public health data the country (29,36). In 2004, the report of snakebite cases became mandatory for each medical center, but later in 2008 the report became individual for each patient (37). The only snakebite incidence report prior 2004 was from 1975 to 1999, with an average of 0.18 envenoming’s per 100.000 inhabitants nationwide (70.8 cases per year) (29). In 2005 SIVIGILA reported 5.03 envenoming’s per 100.000 inhabitants (2161 cases per year), and later, in 2016 the reported incidence was of 9.6 envenoming’s per 100.000 inhabitants (4704 cases per year). This sudden increase of cases seems to be associated with implementing the new reporting system rather than a change in the dynamics of snakebite (36,38).

Nevertheless, the total number of reported cases is still expected to be less than the total cases due to the complexities of data collection in the territory (4,6,18,27,29,39). In Colombia, antivenom cost is borne by the government (through a health provider company), but its distribution usually does not reach the needs areas (12,40). Therefore, the disease burden estimation through mathematical and statistical approaches could help determine the total number of snakebite cases in the country and the real demand for antivenom, thus helping to fulfill the global strategy of the world health organization (10,14,16,41,42).

Our paper focuses on estimating underreporting in the country using a mathematical model that can be easily extrapolated to other countries. To calibrate this model, we first developed a risk estimator based on the distributions of *Bothrops asper* and *B. atrox*, and we tested its performance with snakebite public health records. Second, we used a generalized mixed effect model based on previous work (16), which incorporates the risk estimator as a proxy to estimate total snakebite cases, and poverty data with an accessibility index to estimate the reported fraction of these total cases in each administrative unit. Third, we calibrated our model with Spatio-temporal data of snakebite in Colombia from 2010 to 2019 by using the Markov-Chain-Monte-Carlo (MCMC) algorithm in NIMBLE r-package (43), and finally, we produced maps of underreporting and total incidence that report the real burden of this NTD.

## RESULTS AND DISCUSSION

We constructed a mathematical model which can estimate snakebite incidence underreporting in Colombia at a municipal level. Our estimates are based on a risk map, an accessibility score, and a poverty index. Before exploring the performance and limitations of this model, we want to define the inputs required for its application in a different country: i) A set of incidence data at the finest administrative level, for other months or years, for at least five units of time. Our model estimates underreporting of snakebite incidence given a set of underreported data, but it does not estimate incidence without data. Even so, some modifications can be done to perform a less robust estimation in areas without data, as was discussed by Bravo-Vega et al., (2019); ii) Estimates of distribution for the most relevant venomous snake species, done by performing fieldwork (16), or by performing a niche modeling over a set of presence locations; iii) Human population datasets at a certain administrative level. This data must be in the same temporal and administrative scale as the incidence data; iv) Data of main roads and fluvial routes, which can be generated by using open-source data such as Google Maps or national data from ministries of transport; v) Climatic data, available in open-source servers such as WordClim or TerraClimate (44,45); vi) Geo-referenced locations of every medical center with the capacity to deal with a snakebite; and vii) A poverty estimator at the same spatial resolution than incidence and population data. These data are available for several countries, where the real snakebite burden can be approximated with our approach: This estimation can ultimately help to improve planning strategies and decrease the high threat that this NTD represents to the affected population.

### Risk estimation validity

Our initial snake presence database had 636 records for *Bothrops asper*, and 374 records for *B. atrox*. Our final dataset had 104 and 65 records for each species, respectively. After selecting the best distribution models capturing snakebite risk spatial distribution, the selected model for *B. asper* had a train data AUC of 0.75, a train data CBI of 0.847, and the minimum AICc of all the models. For *B. atrox* the statistics were 0.65 (AUC), 0.76 (CBI), and the difference between its AICc and the model with the minimum AICc was 6.8. Thus, the distribution models generated are adequate to differentiate between suitable and non-suitable areas for both species and significantly better-predicting distribution than a random model(46). Both models suggest that the most suitable areas for *B. asper* are the lowlands of the Middle Magdalena Basin, South Pacific region, and regions of the Caribbean Coast, while the lowest suitability was in the extremely dry areas of the northeast the country (La Guajira department). In the case of *B. atrox* the most suitable areas were located on the lowlands of the Orinoco and Amazonian regions, being suitability relatively constant, while the lowest suitability was on the piedmont of the eastern Andes (View Figure 1).

**Figure 1.**
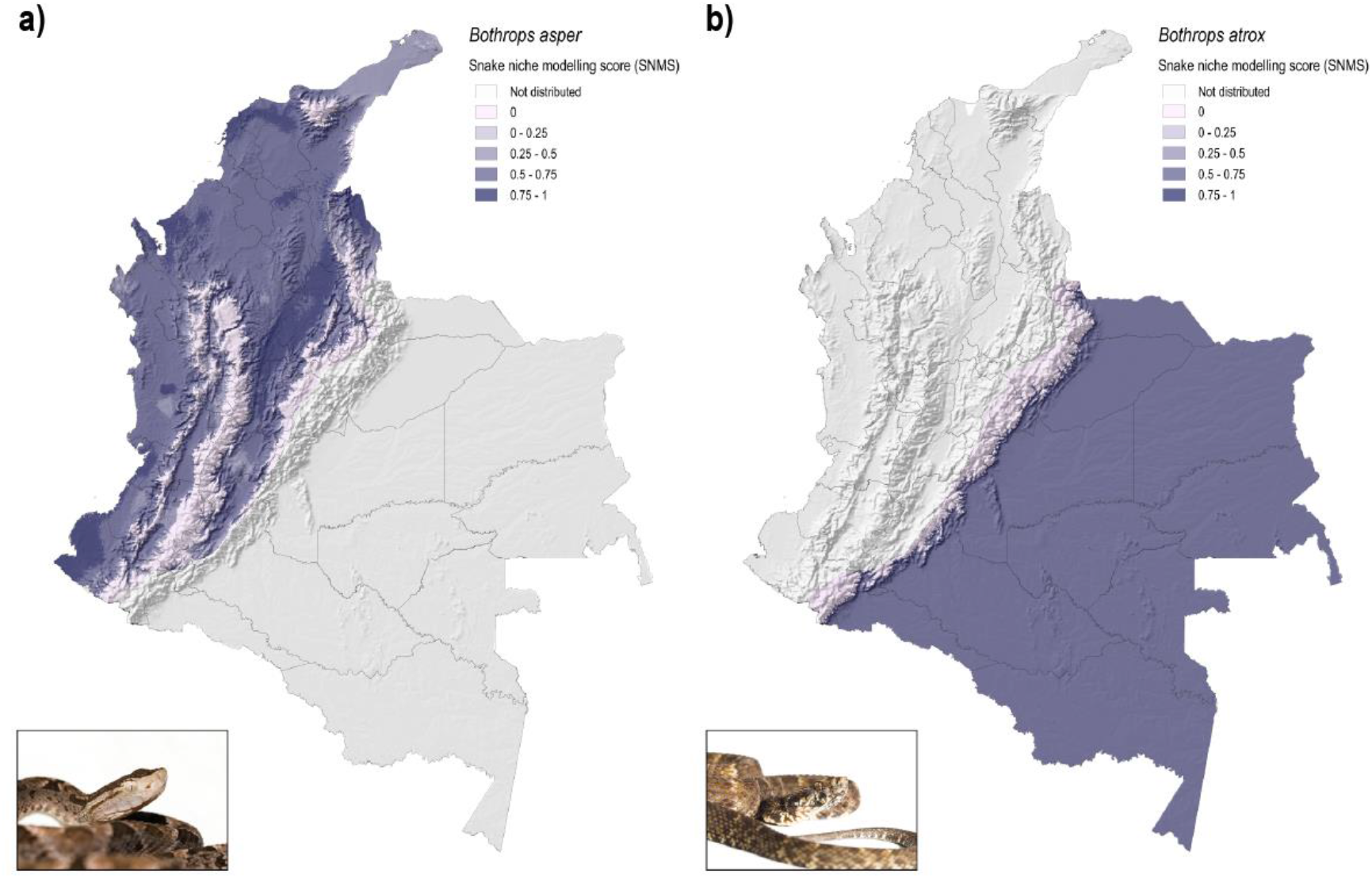
Snake niche modeling score (SNMS). Light purple areas mean 0 niche modeling scores, while light grey areas are outside each species range. Dark purple areas are high suitability areas. Both maps were produced using open-source Geographic Information System Quantum GIS (QGIS). *a) Snake niche modeling score for Bothrops asper*. Suitable areas are in the lowlands of the middle Magdalena basin, south Pacific region, and regions of the Caribbean Coast, while the lowest suitability was in the extremely dry areas of the country’s northeast (la Guajira department). Note the lack of presence locations in the south of the Pacific and in the lowlands located in the south of the Caribbean coast, where is difficult to sample because of social conflict (Figure S1). *b) Snake niche modeling score for B. atrox*. Suitable areas are in the lowlands of the Orinoco and Amazonian basin, while non-suitable areas are located close to the piedmont. Note the low sampling effort in regions of the distribution of the snake, where several pristine and isolated forests are located, and data is absent (Figure S1).

The estimated distribution for both species (Figure 1) seems adequate since our algorithm performance statistics (AUC and BIC) are greater than 0.5, and similar to reported values for successful distribution predictions for different NTDs (19–21,47). In addition, given that we selected the models primarily based on AICc values and then based on the performance capturing spatial variation of snakebite incidence, we can rely on our distribution maps (Figure 2). The maximum entropy algorithm that we used in this study has been applied successfully in predicting the distribution of venomous snakes under climatic change conditions in America (20), and predicting snakebite risk based on distribution estimations of venomous snakes for Veracruz, Mexico (19,20). Other studies have also successfully predicted vector distribution for different zoonotic diseases, as Leishmaniasis and Chagas disease, by using the same algorithm (21,47). The envenoming risk score, which was constructed based on the estimation of the *Bothrops asper* and *B. atrox* distribution models, had a significant linear correlation with the logarithm of the reported risk by SIVIGILA (Pearson’s correlation coefficient: 0.77, p-value < 0.001). Thus, our distribution estimations are valid, and our risk map can be a valuable tool to address the spatial heterogeneity of snakebites in Colombia.

**Figure 2.**
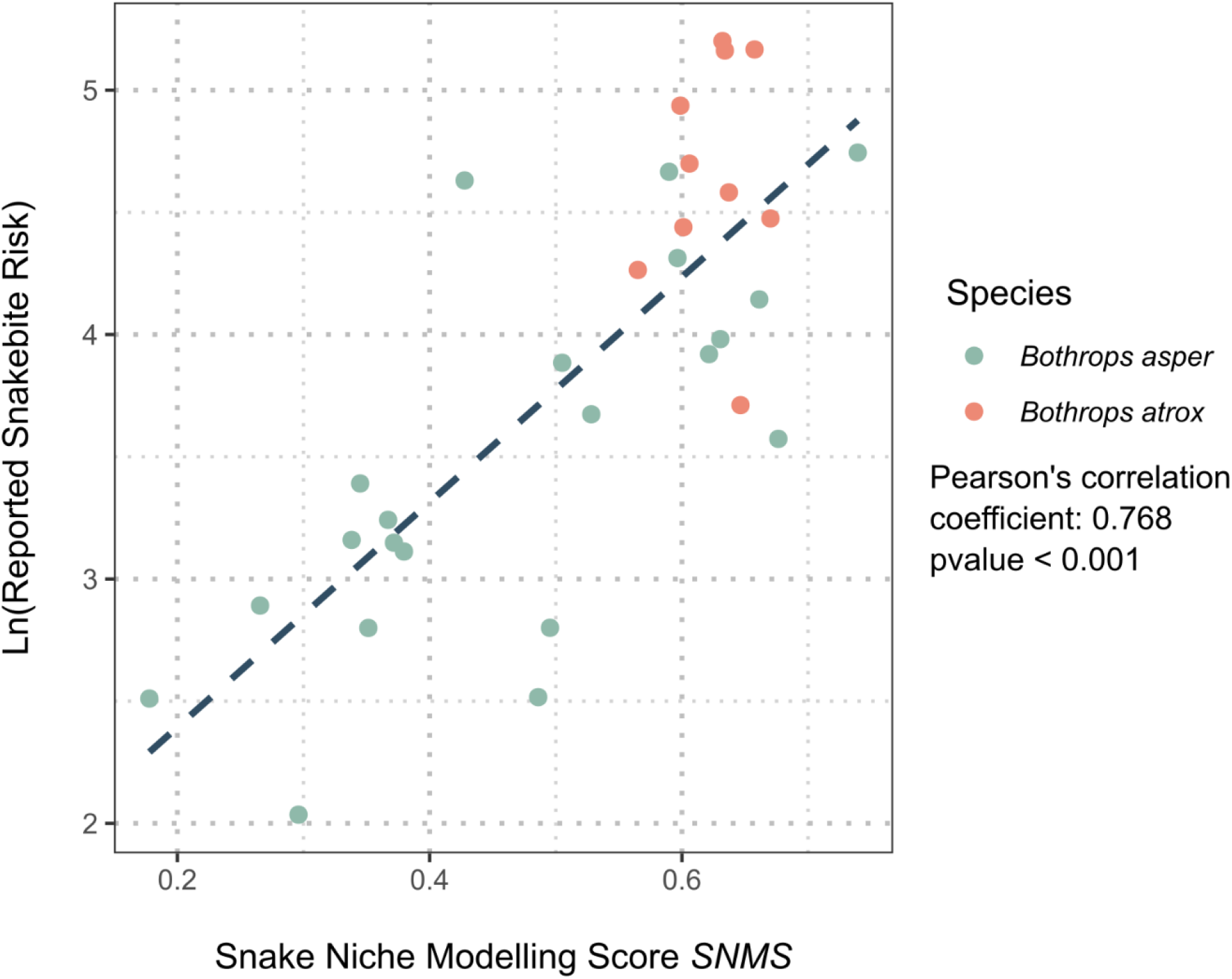
Linear regression between the logarithm of SIVIGLIA (2010-2015) reported risk and envenoming risk score (SNMS) at a departmental scale. Y-axis is the logarithm of SIVIGILA’s reported risk (Envenomings per year per 100.000 persons), and X-axis shows the average envenoming risk score per department. Each point corresponds with a department, and red circles denote departments where *Bothrops asper* is distributed while blue circles are departments where *B. atrox* is found. Note the linear relationship between the two variables, which was quantified with a Pearson’s correlation coefficient of 0.768 (p-value < 0.001). Note that departments with *B. atrox* make a group characterized by high incidence and high envenoming risk score.

We found that both species habitat suitability is higher on humid lowlands, and it is explained not only by total precipitation or temperature but also by their seasonality. These environmental variables have not been used to explain snakebite incidence, where most studies use total precipitation, temperature, and marginalization index as potential incidence predictors (48–52). We want to emphasize the importance of understanding the ecology of venomous snakes to address the environmental factors associated with snakebite.

### Spatial distribution of snakebite risk in Colombia

Most of the country is under high snakebite envenoming risk (Figure 1). Our estimated snakebite envenoming risk is correlated significantly with reported incidence by SIVIGILA (Figure 2), so we can validate our risk estimation as a fine-scale envenoming risk map for the country. When we compared the estimated risk with the reported incidence, we found that higher risk values and reported incidence tend to cluster in departments where *Bothrops atrox* is distributed (Figure 2). Furthermore, these departments are characterized by low population density, low road infrastructure coverage, indigenous populations, and low urbanization (53–55).

We have two not mutually exclusive hypotheses that can explain this higher risk in areas where *Bothrops atrox* is found: i) Socioeconomic disparities can increase the risk of snakebite: Most of the municipalities with lower socioeconomic indicators have low population density, where the most population is rural, having low urbanization(55), which has been demonstrated to be related with snakebite incidence (4,33,37,39,48,56,57). In addition, the intensification of anthropogenic activity (e.g., deforestation and expansion of crops lands) can also increase the risk of snakebite (37,58). Those characteristics could increase the encounter between humans and venomous snakes, explaining the high risk in these areas. ii) Biological factors can increase the risk of snakebite: Given that *B. atrox* and *B. asper* are different species, several biological variables vary among them. For example, it is known that the abundance of *B. atrox* in the Brazilian Amazon is higher close to rivers: Most of the human population in the amazon lives close to water streams because it is the primary connection between towns (59–62). Additionally, the population density could be higher for *B. atrox* than for *B. asper*, and based on studies on feeding ecology *B. atrox* seems to be more active during the day than *B. asper*(63–65). These differences can explain the increase in risk in these departments, but sadly these biological data are not known in Colombia.

We want to stress the importance of collecting natural history data of Colombian venomous snakes to improve the understanding of ecological aspects of snakebite epidemiology. The datasets would allow us to mechanistically understand if the higher risk of snakebite in departments with *Bothrops atrox* is due to socio-economic factors, biological (environmental) factors, or both, to enhance the prevention and control programs to decrease snakebite burden in these risky areas. One way to answer whether biological factors can explain this difference is by performing fieldwork. A previous study in Costa Rica, which is the base of our model, did a census method to measure the relative abundance of *B. asper*, and a mathematical model based on this relative abundance explained the geographic distribution of snakebite cases(16). Performing this methodology in Colombia for *B. asper* and *B. atrox* could clarify differences in the encounter rate between humans and these snakes.

### Underreporting estimation

We evaluated model convergence with the maximum index for the Gelman-Rubin diagnostic which was 1.05 for the parameter *α*_1_, and the multivariate index was 1.02 (66,67), indicating convergence because both numbers were lower than 1.1. Also, prior and posterior distributions for model parameters are shown in Figure S2. We estimated an average number of snakebite cases between 2010 and 2019 of 5227.6 events per year, whereas SIVIGILA only reported an average of 4689.3 yearly events for the same time. As a result, we estimated that 538.02 cases are not reported each year, corresponding to 10.3% of the total cases. The geographic distribution of reported cases and our estimation of total cases at a municipality scale can be seen in Figure 3. Most of the envenoming reports were from Antioquia, having around 683.4 cases each year (Figure 3.a). Thus, each year the number of cases that do not get medical attention in the country is similar to the reported cases of the most affected department (National not-reported cases correspond with 78.7% of the cases that Antioquia reports). Interestingly, departments where *Bothrops asper* is present report most snakebite cases, but the departments with *B. atrox* are riskier (Figure 3). This pattern corresponds with results observed in Figure 2, where we see that in areas where *B. asper* is distributed, more cases occur because of the high density of human population in Colombia (68), but in areas where *B. atrox* is distributed people are more prone to suffer a snakebite.

**Figure 3.**
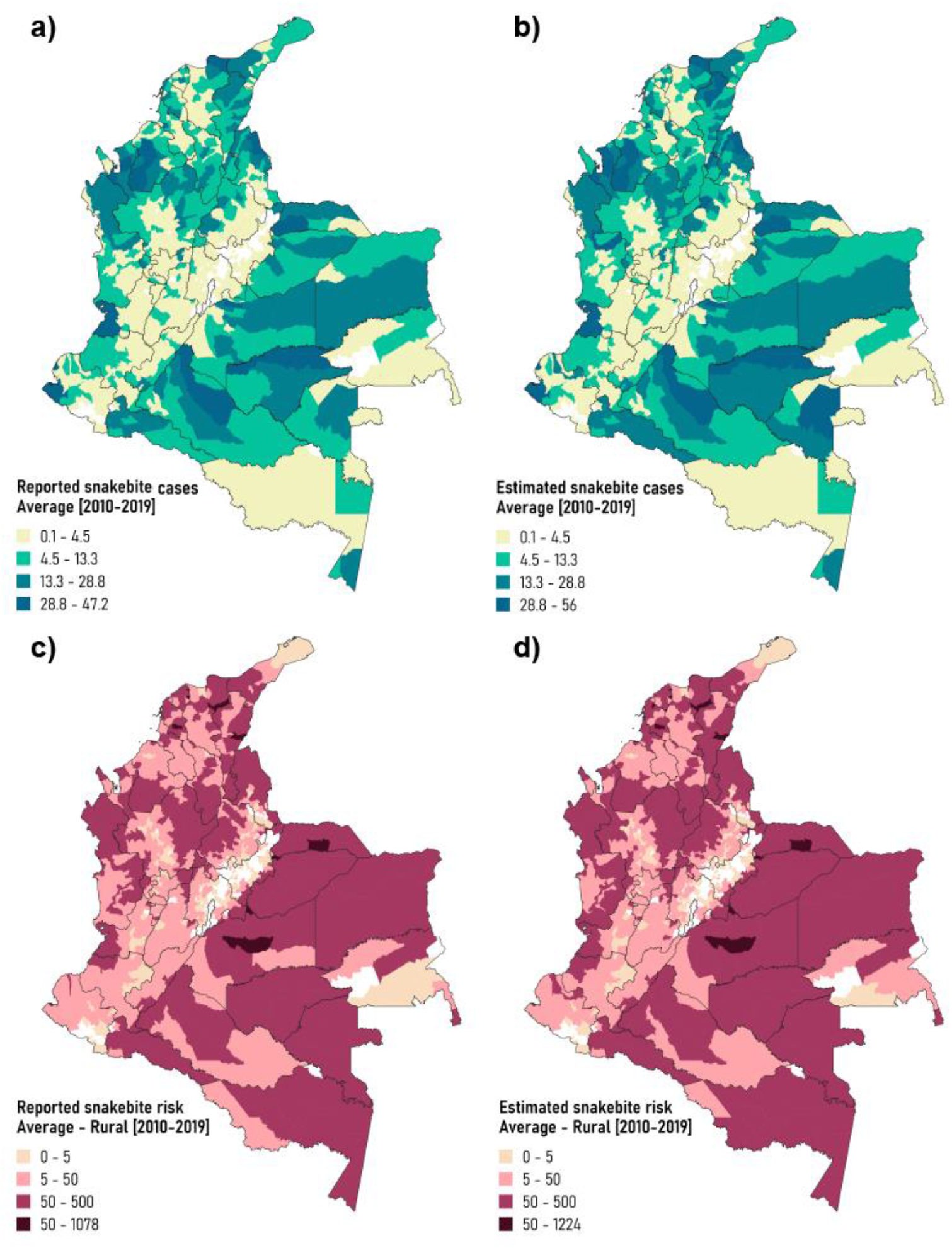
Reported and estimated snakebite incidence and risk at a municipality scale. Estimates were done with the calibrated model after parameters estimation by MCMC. a) Yearly reported SIVIGILA cases (2010-2019). Note that the spatial distribution of the cases is heterogeneous, where the department that reports most envenoming’s is in Antioquia. b) Yearly estimated total cases. Note that most municipalities increased in the scale of cases, indicating that underreporting affects most of the country. c) Yearly reported SIVIGILA snakebite risk. Areas with higher risk are located in the Orinoco and Amazonian areas, where *Bothrops atrox* is distributed. In addition, the riskiest departments are Casanare, Guaviare and Vaupes. d) Yearly estimated total snakebite risk. Most of the country have more risk than the reported ones, where departments as Amazonas and Arauca are now located in the highest risk group of departments. On the other hand, departments with *B. asper* as Cesar, Atlantico, and Norte de Santander are now the riskiest departments with this species.

Mortality rate in a population that do not receive medical attention can increase severely (2,5,6,69–71). A report from African venomous snakes estimates that the reduction of lethality caused by antivenom was from 10-20% to 2.8%. This is the only report of lethality reduction available in the literature (72). Then the cases that are not reported because individuals did not seek medical attention will be more prone to be fatal, increasing the underreporting fraction drastically in deaths caused by snakebite. In addition, according to the distance to the nearest medical center map (Figure S3), we estimated that 68% of the country is located at more than two hours to the nearest medical center, while 36% of the country is more than 12 hours faraway from these centers. The country’s low coverage and accessibility to public health increase the time between the envenomation and the medical attention. Most country areas exhibit large values of these factors, resulting in severe envenoming’s with an increased risk of disability or mortality (3,70). Colombian health authorities have made a considerable effort to enhance the reporting system, and now we have a strong data gathering system compared with other countries (37), However, there is still a considerable underestimation (View Figure 3 and Figure 4 that shows that underreporting affects the whole country): Snakebite is a critical life-threatening public health issue that is still neglected by authorities in tropical countries, and its real threat to human life is still not quantified.

**Figure 4.**
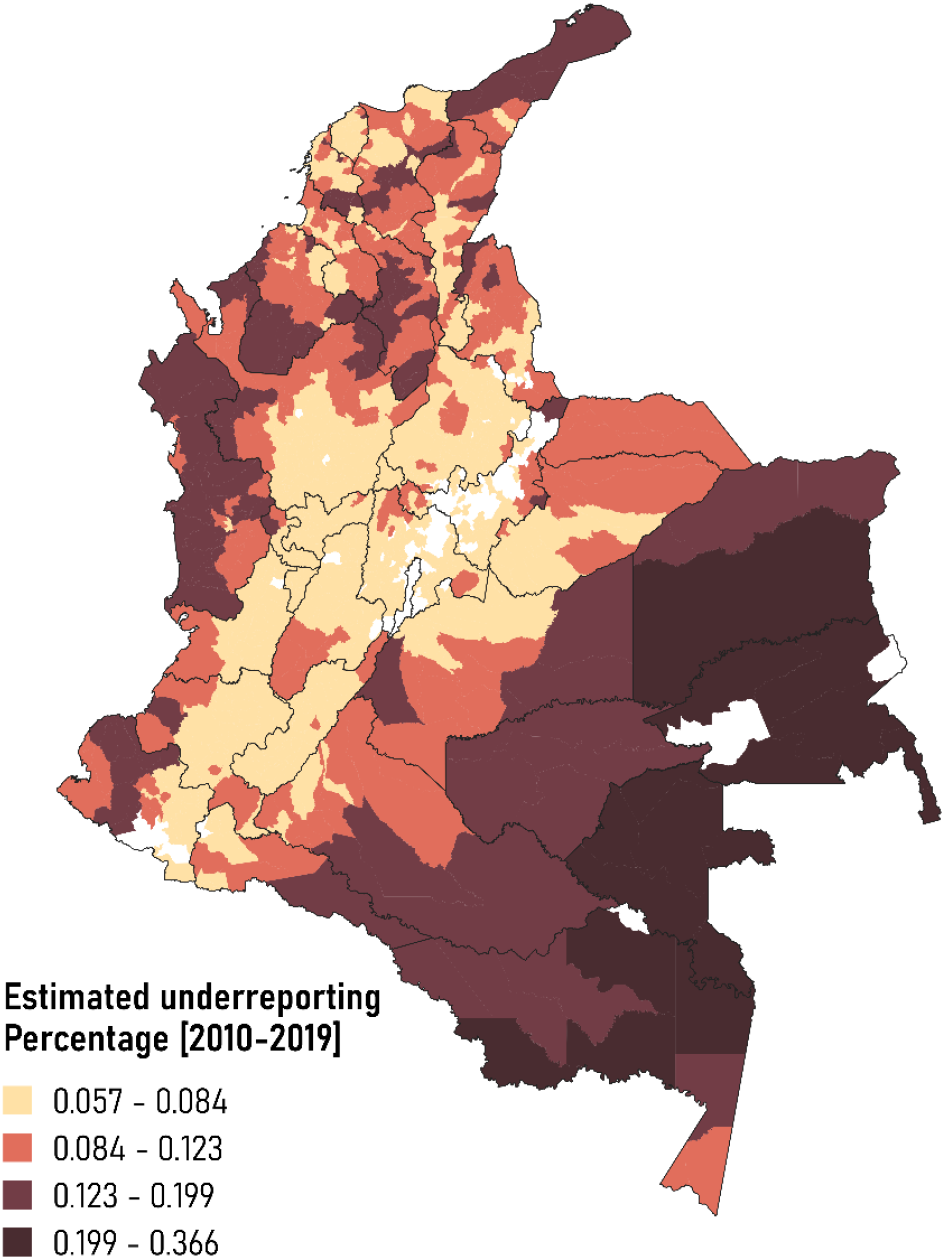
Estimated snakebite underreporting. The areas most affected by underreporting are the southeastern part of Colombia, where *Bothrops atrox* is distributed. Note that the pacific and northern Caribbean coast region also suffers from underreporting.

Our under-reporting estimates vary between 5.7% and 38.7% of total cases, which indicates that underreporting is a relevant problem that affects data collection in snakebites in the country (Figure S4 shows the relationship between underreporting, travel time, and unsatisfied basic needs after model calibration). In addition, areas located at the eastern and southern part of the Amazonian and Orinoco regions are the most affected by underreporting, sharing the highest snakebite risk (Figure 3.c-d). Moreover, the Pacific and northern Caribbean coasts also have a high underreporting, where these areas share a high poverty index and the longest travel time because of poor vial infrastructure (4,39,42,73–75). An underestimation in the reported cases due to lack of accessibility could increase snakebite mortality and morbidity. Furthermore, the treatment by antivenom is not distributed evenly, causing that places with low reporting rates have no antivenom to attend the envenomings (2,6,69–71). Thus, our underreporting estimation can be used to enhance antivenom availability and rural population inclusion into medical insurance (12,42,76–78).

### Limitations

Although our proposed model can estimate underreporting in snakebite incidence using modern mathematical tools, we want to discuss a few limitations that can be improved to obtain more reliable estimations. First, we believe that our risk estimation based on snake distributions is a significant contribution, but environmental niche modeling is limited by the quality of the species presence records. For example, several areas in Colombia and worldwide have limited accessibility, and thus biological data may be scarce (View Figure S1) (79). Nevertheless, the maximum entropy algorithm has shown good performance in estimating distributions for species with incomplete presence data (80–83).

Second, the assumption that the exposed population is the total rural population may not be accurate. Several municipalities have broad elevation gradients, where rural areas are located in a high elevation where no venomous snakes are present. Additionally, extreme urbanization would decrease the abundance of venomous snakes; hence, people living in areas under this condition will not be as exposed as the rural population but will suffer snakebite (84–86). Even so, our risk score is an reliable estimator for snakebite risk in Colombia based on the significant correlation between the reported incidence at a departmental scale and our envenoming risk score (**¡Error! No se encuentra el origen de la referencia**.).

Finally, we assumed that accessibility to treatment is only dependent on the distance to the nearest medical center (Figure S3). Other factors affecting accessibility to treatment are the correct training to medical personnel and the optimal distribution of antivenom (9,13). We assumed that real snakebite incidence only depends on our proposed risk estimator, which is based on snakes’ occurrence, but it could be also mediated by human activities (17,87). We also assumed that underreporting only depends on the accessibility score and a poverty index, but it could depend on other cultural, economic, and public health logistic factors (18,88,89). Our model can estimate based on available data and a strong and reliable statistical method. This framework is one of the most current tools in model fitting, and this study is the first time these tools are applied to snakebite. So we suggest it as a reliable tool to perform the most educated guess about snakebite underreporting, which can improve disease management but will never replace a correct epidemiological data gathering, which is the responsibility of national public health instances.

### Final remarks

Using geolocations of venomous species and available environmental information, we could estimate the distribution of the two most important venomous species in Colombia, which explains the geographical variation of reported snakebite incidence in Colombia. Furthermore, we believe that new insights about snakebite can be obtained by including natural history traits of the venomous snakes to understand its epidemiology (9). For example, different studies for *Bothrops asper* in Costa Rica and *B. atrox* in Brazil looked at the trophic ecology of these species and its relationship with their habitat usage, where their findings can help determine risky micro-habitat conditions which can be targeted in order to prevent snakebite (63,64,90). On the other hand, reproduction studies for *B. asper* in Costa Rica highlighted the relationship between the birth seasons of this species and the seasonality of incidence, so the temporal patterns for this event were established (49,91). Sadly, in Colombia, information about the ecology of venomous snakes is scarce or non-existent (34). In addition, we developed an underreporting estimation by using advanced inference, mathematical and statistical tools, and available biological data: Even so, the reporting system in Colombia is one of the strongest in the area (37), this study is critical to quantify the real burden of snakebite and to give a framework that can be used in other countries with a data-reporting deficit. Although our modeling approach has several strengths, we want to stress that this framework will never substitute correct surveillance that must be the responsibility of public health instances.

Finally, we would like to suggest five strategies to improve the management of snakebite in Colombia: i) To understand how micro-habitats affect the presence of venomous snakes close to domestic areas: Our proposed species distribution maps for this study have fine-scale but are still coarser enough to capture microclimate household variation, which affects the encounter frequency between a venomous snake and a human. ii) To determine if snakebite envenoming has seasonality in this country and if it is related to environmental seasonal patterns: We found spatial environmental variables which correlate with the geographic distribution of snakebite incidence, but we still do not know how these variables are mechanistically associated with the dynamics of snakebite envenoming. iii) To improve the distribution strategies for antivenom: Our model can help determine areas in need of antivenom availability and estimate the total number of vials needed to satisfy the demand of the treatment. Then, the optimal distribution of this valuable resource will reduce the burden of this Neglected Tropical Disease overexposed populations. iv) To improve the capacities of health staff in the management of envenoming: The correct application of the antivenom is crucial to decrease the venom effects, so national programs that aim to train medical staff to deal with a snakebite will improve the effectiveness of the treatment. v) To include snakebite into the public health scope to strengthen resources and to improve the empowering of populations in public health processes: This is aligned with the global strategy developed by the WHO (14), where the results of this study increase the visibility of the problem, but snakebite is still not included into national public health scope. Finally, snakebite envenoming fulfills all the characteristics related to a Neglected Tropical Disease, and its burden in Colombia is high: It is time to include this disease into the national public health scope.

## METHODOLOGY

To generate and calibrate a generalized mixed-effect model that estimates snakebite incidence underreporting, we followed three steps: First, we built an envenoming risk score map based on snake distributions (obtained through ecological niche modeling) and the law of mass action. Second, we calculated an accessibility score that reflects the struggle to reach a medical center. Finally, we developed and parametrized a mixed mathematical model to estimate underreporting.

### Snakebite’s envenoming risk score map

The envenoming snakebite risk (Defined as the probability that envenoming will occur in a determined population during a time) is mainly associated with the presence of venomous snakes. Therefore, we developed an envenoming risk score incorporating the ecological niche modeling scores for the two species (*SNMS*, see below), using snakes and environmental layers as input presence records. This risk model is based on the law of mass action, which states that envenoming will be the result of encounters between humans and snakes, and it will be proportional to the multiplication of their abundances (16):

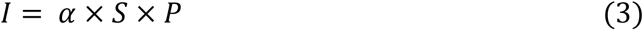

In Eq. 3, *I* is the snakebite incidence, *S* is the snake’s abundance, *P* is the rural population, and finally the contact rate between both populations. Dividing the equation by *P*, approximating *S* to *SNMS*, and using a logarithmic transformation on the reported risk to linearize its relationship with *SNMS*, we will obtain our estimation of risk (Eq. 4):

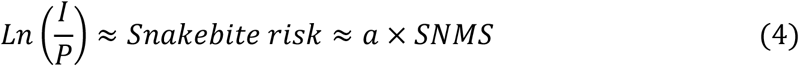

#### Snake presence data

We selected *Bothrops asper* and *B. atrox* as study species because of their wide range, ability to adapt to human intervention, and responsibility for most snakebites in Latin America and Colombia (25,27–29,32,33,35,64,92). First, we built a database of presence records using georeferenced samples obtained from natural history museums in Colombia (See acknowledgments), where duplicated records were eliminated for each species. Then, to homogenize the spatial data, we used a grid of 0.3° resolution for *B. asper* and 0.7° resolution for *B. atrox* to select only one presence point per square-grid (View Figure S1). Finally, we removed records above 1900 m.a.s.l. for *B. asper* and 1500 m.a.s.l. for *B. atrox* because these are the maximum altitude thresholds known for their distribution in South America (25,31). Our initial snake presence database had a total of 636 records for *B. asper*, and 374 records for *B. atrox*. After data depuration, our final dataset had 104 and 65 records for each species respectively.

#### Environmental layers

We used Bio-climate variables from the WorldClim database server (http://www.worldclim.org) (44) for the current conditions in a resolution of approximately 1 km x 1 km. We removed collinear environmental layers with the package *Virtual Species* in R (93,94). We used a threshold based on Pearson’s R correlation coefficient of 0.9 (95) to subset non-correlated environmental layers. The selected Bioclimatic variables are shown in Table S5.

#### Envenoming risk map estimation and validation

We predicted species habitat suitability using a maximum entropy algorithm because we used only presence data (81). After performing the algorithm described in S6 to generate a set of maps representing the ten best distribution models for each species, we generated species distribution predictions from these best models over each species range to produce habitat suitability maps using a cloglog output format (96). Then, we removed areas above known altitude thresholds for both species (25,31). Finally, given that both species are not sympatric (25,31), we combined all the possible permutations of these ten predictions to obtain 100 maps of both species habitat suitability in Colombia. These score maps are the snake niche modeling score per pixel (*SNMS\**), with the same resolution as the environmental layers.

To select the best *SNMS\** map to be proposed as our final envenoming risk score map (*SNMS*), we computed the average value of *SNMS\** for each department for each one of the 100 generated maps, and we compared it with the reported snakebite risk (snakebite incidence per 100.000 persons). To compute reported risk, we averaged SIVIGILA reported incidence between 2010 and 2019, and we obtained rural population estimates for the same timespan from the National Administrative Department of Statistics (DANE). SIVIGILA’ reported incidence corresponds to the cumulated cases for each year for each department (36), and it reports the number of people that received medical attention after suffering a snakebite.

To compare *SNMS\** with reported risk, we used a type II linear regression between the 100 *SNMS\** maps and the logarithm of reported risk (View Eq. 4) because both variables are random variables, and we estimated regression coefficients by using ordinary least squares(97). Finally, we selected the *SNMS\** map with the highest correlation with reported risk as our final envenoming risk score map (*SNMS*), a valid snakebite risk estimator. Maps were constructed using the open-source Geographical Information System Quantum GIS (QGIS).

### Computation of the accessibility score

To compute the accessibility score, we assumed the distance mainly determines it to the nearest health center. For this, we used a georeferenced dataset of clinics, hospitals, and health centers in the 2016 Geostatistical National Framework published by the DANE (68). Distance to a medical center is essential in determining underreporting because snakebite mortality is mainly associated with the difficulty of having access to medical attention on time, so people who live far away from medical centers will seek another kind of treatment (56). Other aspects that can contribute to accessibility are the availability of serum or the presence of trained personnel (33,56,98). Unfortunately, we did not have access to information about these factors, so we decided to compute our score using only distance as a first step towards understanding the effect of antivenom availability on underreporting. Hence, we first created a combined map of travel speed by using maps of: i) Roads and type of road from the World Food Programme and Open Street maps (99), ii) Land coverage from the Food and Agriculture Organization of the United Nations (100), iii) Fluvial transport from the Agustin Codazzi institute (101) and *iv)* Geographic slope map computed using *raster* package in R environment (93,102) and an altitude map from the global server WorldClim (44). Then, we computed the minimum travel time for each location to the nearest medical center based on the travel speed map obtained by merging input maps. This algorithm was performed using the package gDistance in R (103), and the output is a map of minimum travel time to the nearest medical center, which we will assume as the accessibility score.

### Statistical modelling of snakebite underreporting

#### Model Description

The first step of the theoretical-statistical model is to model the *real* incidence of snakebite. As it was shown in (16), the law of mass action correctly represents the real incidence of snakebite. By using Eq. 4, and writing the model as a generalized lineal model, we can state the following:

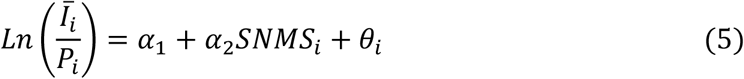

Where *Ī* is the real incidence of snakebite, *α*_1_ and *α*_2_ are the intersect and the slope of the generalized lineal model respectively, θ accounts for a normal random noise, and sub-index *i* denotes the geographical region, in our case municipality, of the underreporting estimate. Assuming the population as an offset we can re-write the model as (Eq. 6):

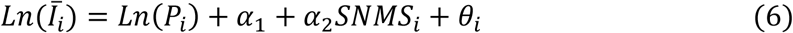

Then, we defined the reporting of snakebite incidence as a counting Poisson process, where the mean is the real incidence (Eq. 6) times a reporting fraction *π* _*i*_. So, the model that will be used to represent reported snakebite incidence *I*^r^_*i*_ is the following:

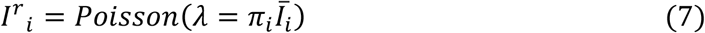

To estimate the reporting fraction we assumed that it will depend on the proposed accessibility score, and on a poverty index (2,13,39,49,56,89). We used the unsatisfied basic needs index as the poverty index because it represents the poverty at a municipality and departmental scale, and we defined the accessibility score for each municipality as the geographic-averaged travel time (75). By assuming a general linear dependence between reporting, the model for reporting fraction is the following (Eq. 8):

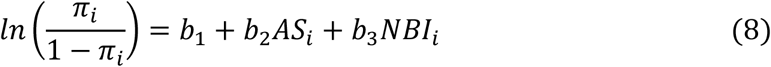

We applied the logit function to the reporting fraction *π* _*i*_ to guarantee that this fraction will be between 0 and 1. In this part of the model, *b*_1_, *b*_2_, and *b*_3_ are parameters, *AS* is the accessibility score, and *NBI* is the poverty index. Thus, the complete model to estimate reported incidence is the following (Eq. 9):

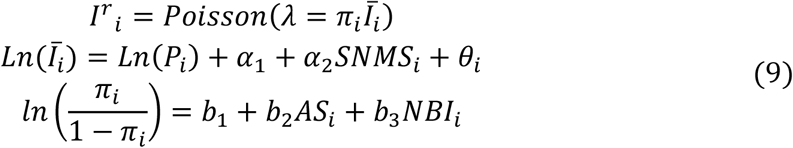

#### Underreporting estimation

To estimate the model parameters we used the MCMC algorithm, which uses a Bayesian approach to fit the model to data (43). This algorithm uses a prior estimation of the distribution of the parameters to finally find a posterior distribution, which maximizes the likelihood between the data and the output from the model. To perform this, it uses the Monte Carlo algorithm to sample parameters from the prior distributions and to compute the likelihood (To see prior distribution view Figure S2). Finally, optimizing this likelihood uses a Markov Chain to iterate the parameter selection and converge to an optimal solution that maximizes the likelihood (104). First, the posterior distributions for the parameters were estimated by using the NIMBLE package in R environment (43,93). Next, we used an automated factor slice sampler (AFSS) to sample the parameters, and then we used 4 chains from different initial conditions to ensure a global optimal for the convergence. We did a total of 4.7 million iterations, where the first 57% were discarded as a burn-in, and we saved the following 43% iterations to check convergence and perform underreporting estimations with the model. To ensure convergence, we calculated the Gelman-Rubin diagnostic by defining a threshold of non-convergence for this index at 1.1, where values above this threshold indicate non-convergence (66,67). Incidence data was obtained from yearly municipality reports of snakebite done by SIVIGILA between 2010 and 2019.

## ACKNOWLEDGMENTS

We thank to the natural history museums of the Universidad Industrial de Santander (UIS), Universidad de los Andes, Institución Universitaria ITM, Universidad del Tolima, Universidad del Valle, Universidad de la Salle (MLS), Instituto Nacional de Salud (INS), Universidad del Magdalena and Universidad Javeriana. Also, to all collections curators who gave access to specimen records, especially Martha Calderón-Espinosa and John D. Lynch (Universidad Nacional de Colombia -ICN), Andres R. Acosta-Galvis (Research institute von Alexander von Humboldt - IAvH), Fernando Sarmiento-Parra and Julieth S. Cardenas-Hincapie (MLS), Martha Patricia Ramírez and Elson Meneses-Pelayo (UIS), Juan Manuel Daza (Universidad de Antioquia - MHUA), as well as all colleagues who kindly share with us their unpublished specimen records, especially Juan José Torres (INS) and Sergio Cubides Cubillos (Butantan Institute). Finally, we thank to Minciencias 727 application for doctoral students during 2016.

## SUPPLEMENTARY INFORMATION

**Figure S1.**
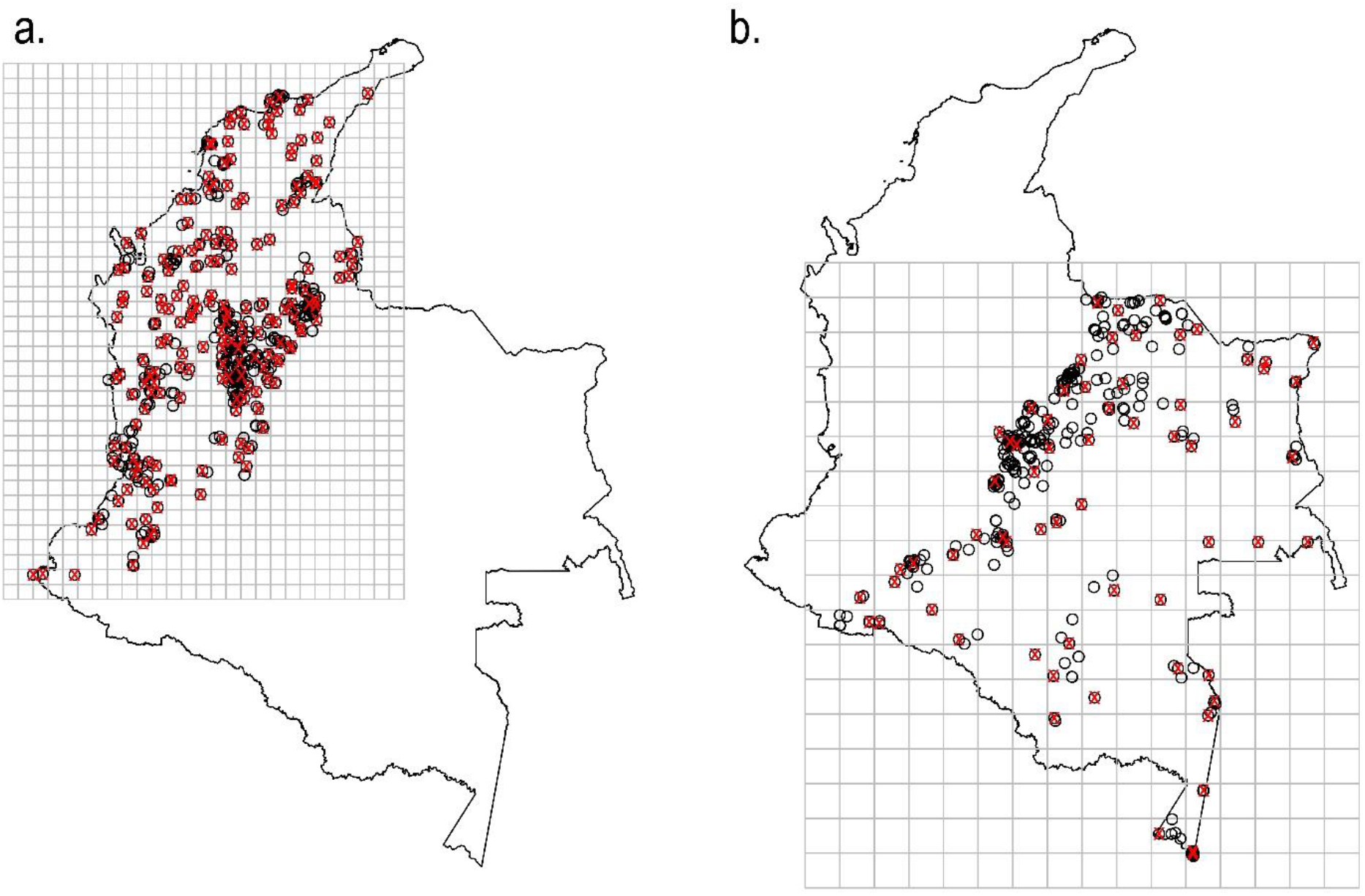
Snakes’ presence records the homogenization process. *a)* Shows the grid used for presence records for *Bothrops asper*. This grid resolution is 0.3°, and we have selected one presence record for each cell in the grid. *b)* Shows the grid used for presence records for *Bothrops atrox*, with a grid resolution of 0.7°, and for each square, we only selected one presence record.

**Figure S2.**
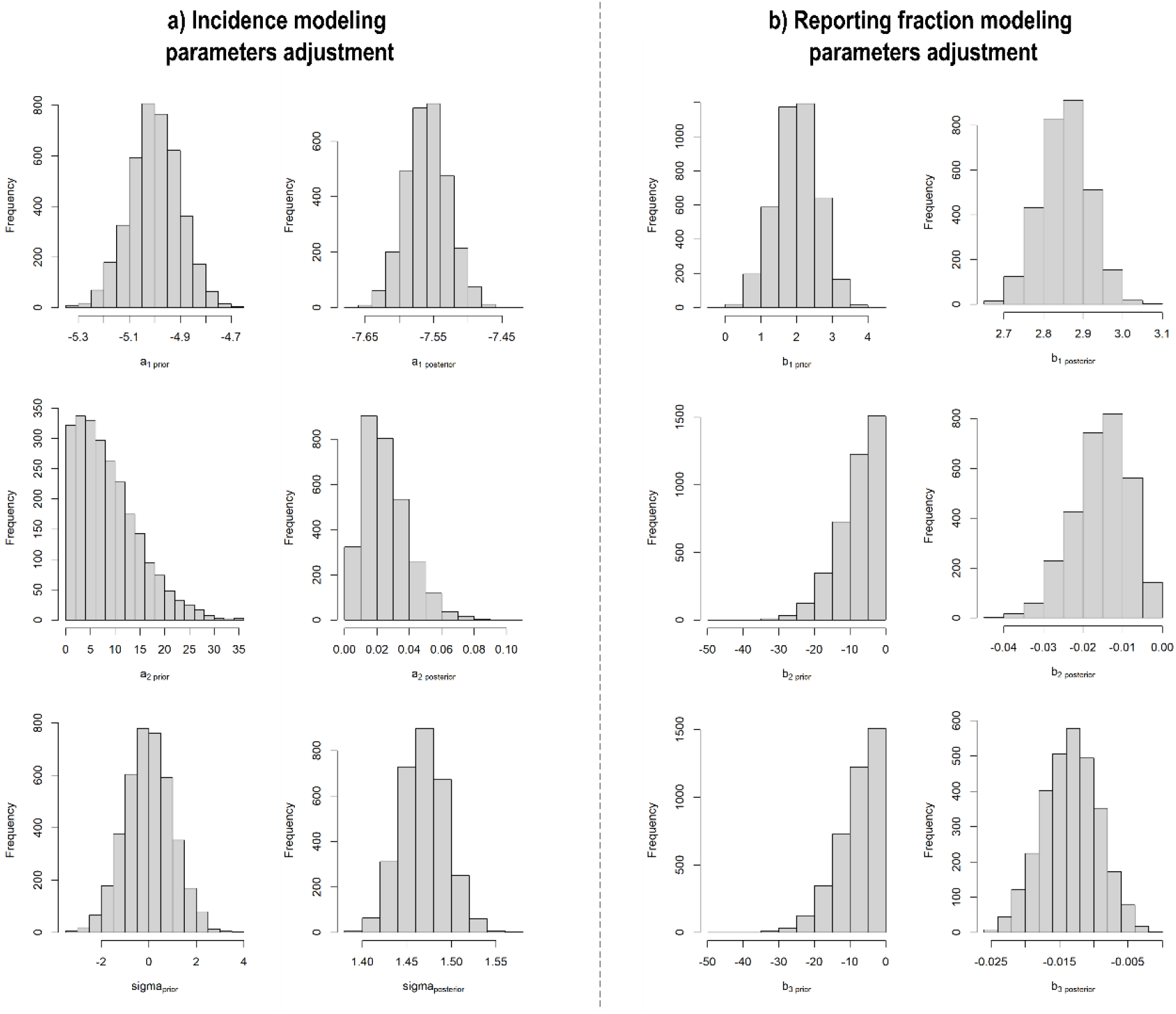
The prior and posterior distribution for model parameters after fitting convergence.

**Figure S3.**
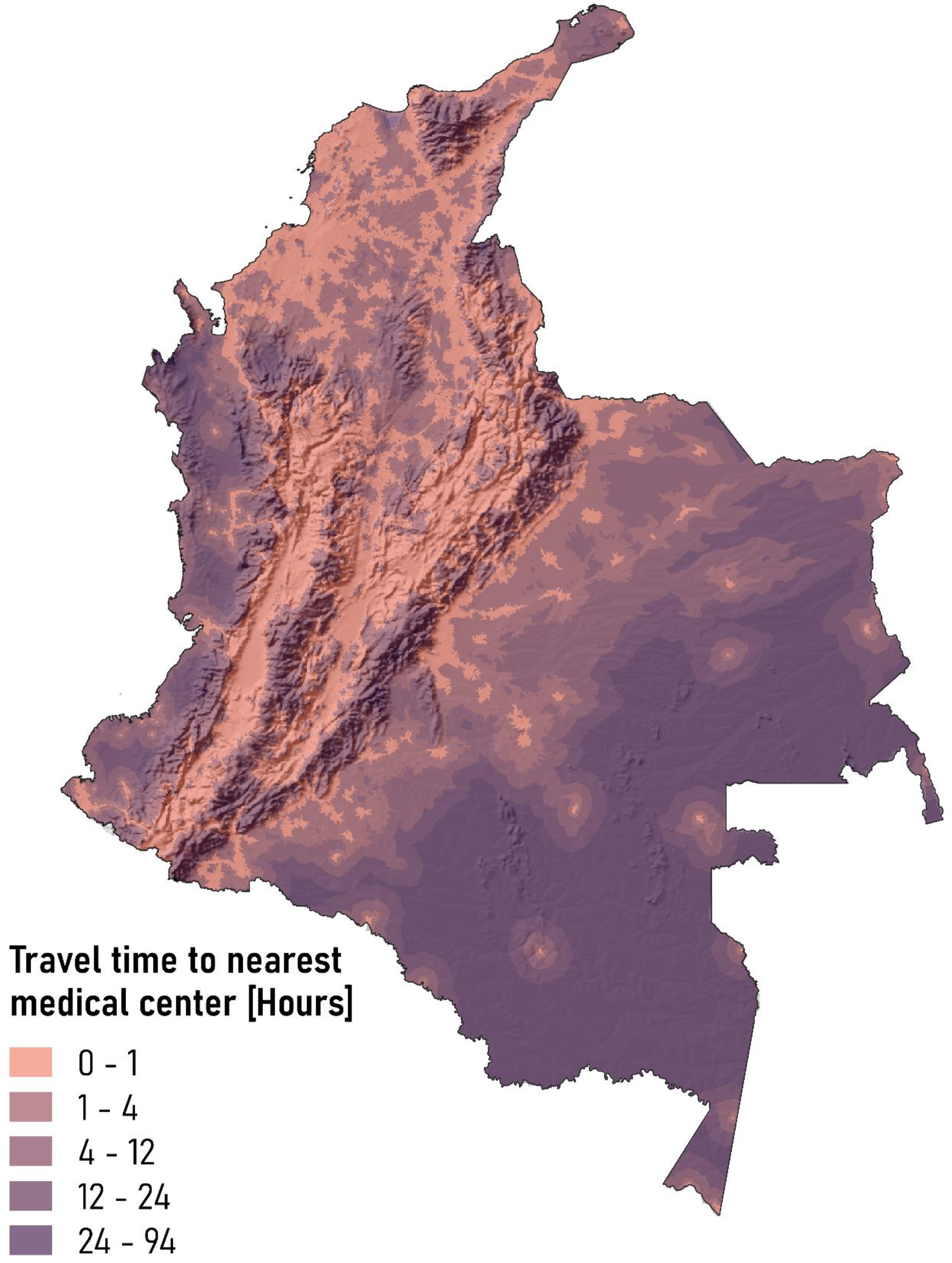
Travel time to the nearest medical center. We used georeferenced data of medical centers with the capacity to administrate antivenom, and we determined travel speed with maps of roads, fluvial routes, slope, and land coverage.

**Figure S4.**
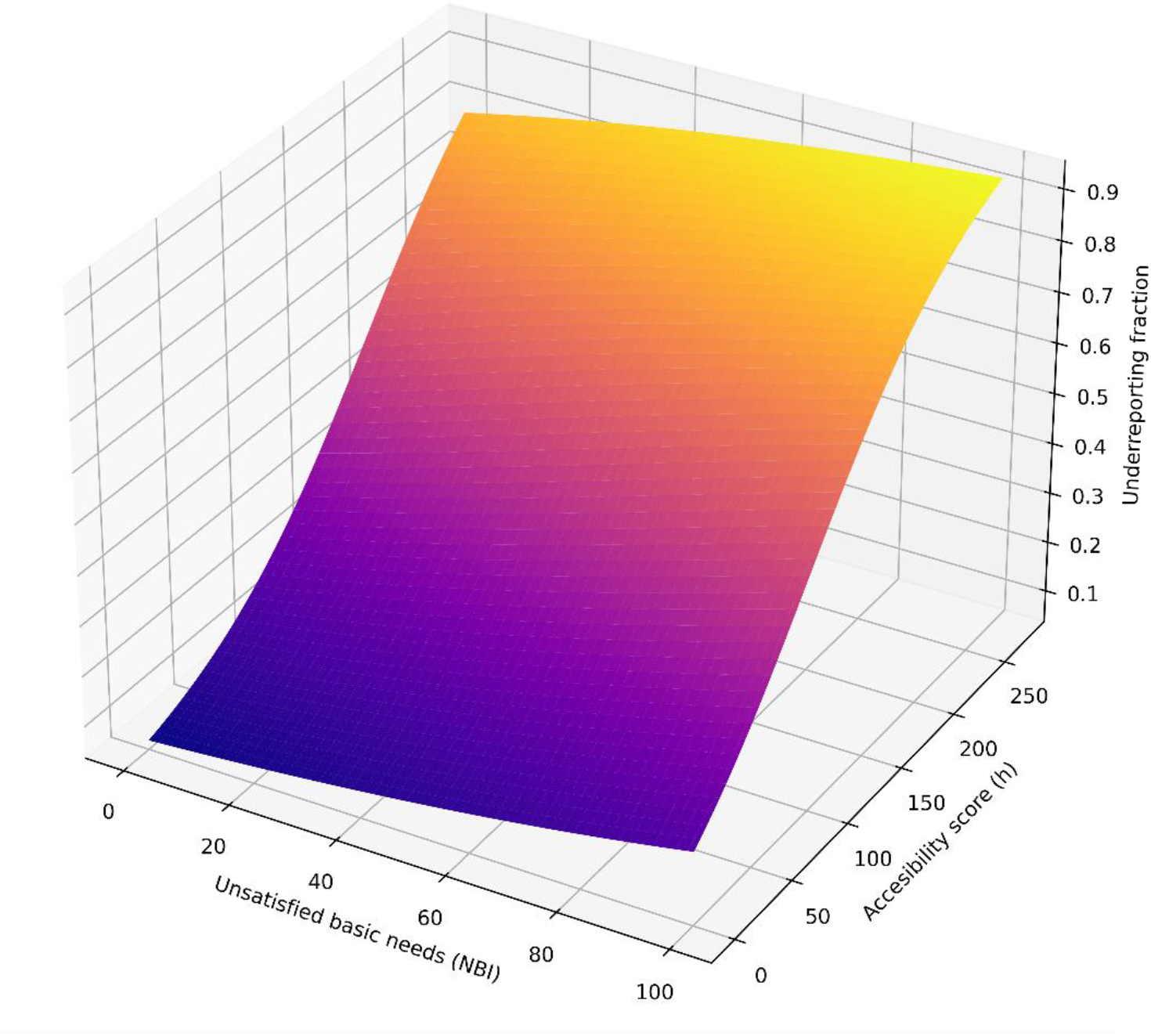
Dependence of underreporting on accessibility score and poverty after model parametrization. As we stated in the model, there is a positive correlation between both variables and underreporting, where low public health coverage and high poverty will increase underreporting fraction. Therefore, our estimation of underreporting varies between 5.7% and 38.7%.

**Table S5.** Environmental layers used for ecological niche modeling.

| Variable | Units | Source |
| --- | --- | --- |
| Annual Mean Temperature | °C*10 | WorldClim |
| Mean Diurnal Range | °C*10 | WorldClim |
| Isothermality | - | WorldClim |
| Temperature Seasonality | °C/100 | WorldClim |
| Temperature Annual Range | °C*10 | WorldClim |
| Annual Precipitation | mm | WorldClim |
| Precipitation of Wettest Month | mm | WorldClim |
| Precipitation Seasonality | - | WorldClim |
| Precipitation of Driest Quarter | mm | WorldClim |
| Precipitation of Warmest Quarter | mm | WorldClim |
| Precipitation of Coldest Quarter | mm | WorldClim |

**S6. Maximum entropy model calibration and selection of the best distribution models**

We used the package *ENMeva*l in r to perform the niche modeling by entropy maximization. First, we defined as background a buffer around the depurated occurrence locations for each species. This buffer had a ratio of 1.5º (Around 166km near the equator), and we randomly selected 100 points over this buffer as background records. This step aims to compare the climatic conditions of each presence location with the climatic conditions of its surroundings. Then, we evaluated different models in *ENMeval* by varying feature classes (These are the potential mathematic shape of the response curves between suitability and climatic variable) and the regularization multiplier (This parameter allows to penalize over-complexed models). We selected the six most used feature classes (Lineal, Lineal-Quadratic, Hinge, Lineal-Quadratic-Hinge, Lineal-Quadratic-Hinge-Product and Lineal-Quadratic-Hinge-Product-Threshold), and also we defined the regularization multiplier between 1 and 6 (81). To partition data avoiding possible spatial autocorrelation in presence records, we used the block partitioning algorithm, which selects 4 blocks in the geographic area that contains approximately the same quantity of presence records, and partition data in each block in 4 points (105). Finally, after performing maxent modelling in *ENMeval* package for each one of the combinations between feature classes and regularization multiplier, we selected per each species the 10 models with the lowest AICc. Our habitat suitability model and its performance evaluation was performed using *dismo* and *ENMeval* packages in *R* environment (93,106,107).

